# *In vivo* microstructural heterogeneity of white matter lesions in Alzheimer’s disease using tissue compositional analysis of diffusion MRI data

**DOI:** 10.1101/623124

**Authors:** Remika Mito, Thijs Dhollander, Ying Xia, David Raffelt, Olivier Salvado, Leonid Churilov, Christopher C Rowe, Amy Brodtmann, Victor L Villemagne, Alan Connelly

## Abstract

White matter hyperintensities (WMH) are commonly observed in elderly individuals, and are typically more prevalent in Alzheimer’s disease subjects than in healthy subjects. These lesions can be identified on fluid attenuated inversion recovery (FLAIR) MRI, on which they are hyperintense compared to their surroundings. These MRI-visible lesions appear homogeneously hyperintense despite known heterogeneity in their pathological underpinnings, and are commonly regarded as surrogate markers of small vessel disease in *in vivo* studies. Consequently, the extent to which these lesions contribute to Alzheimer’s disease remains unclear, likely due to the somewhat limited way in which these lesions are assessed *in vivo*. Diffusion MRI is sensitive to white matter microstructure, and might thus be used to investigate microstructural changes within WMH. In this study, we applied a method called single-shell 3-tissue constrained spherical deconvolution, which models white matter microstructure while also accounting for other tissue compartments, to investigate WMH *in vivo*. Diffusion MRI data and FLAIR images were obtained from Alzheimer’s disease (*n* = 48) and healthy elderly control (*n* = 94) subjects from the Australian Imaging, Biomarkers and Lifestyle study of ageing. WMH were automatically segmented and classified as periventricular or deep lesions from FLAIR images based on their continuity with the lateral ventricles, and the 3-tissue profile of different classes of WMH was characterised by three metrics, which together characterised the relative tissue profile in terms of the white matter-, grey matter-, and fluid-like characteristics of the diffusion signal. Our findings revealed that periventricular and deep lesion classes could be distinguished from one another, and from normal-appearing white matter based on their 3-tissue profile, with substantially higher free water content in periventricular lesions than deep. Given the higher lesion load of periventricular lesions in Alzheimer’s disease patients, the 3-tissue profile of these WMH could be interpreted as reflecting the more deleterious pathological underpinnings that are associated with disease. However, when alternatively classifying lesion sub-regions in terms of distance contours from the ventricles to account for potential heterogeneity within confluent lesions, we found that the highest fluid content was present in lesion areas most proximal to the ventricles, which were common to both Alzheimer’s disease subjects and healthy controls. We argue that whatever classification scheme is used when investigating WMH, failure to account for heterogeneity within lesions may result in classification-scheme dependent conclusions. Future studies of WMH in Alzheimer’s Disease would benefit from inclusion of microstructural information when characterising lesions.

## 1 Introduction

Alzheimer’s disease has long been considered a pathologically-defined disease, whose clinical symptomatology is underpinned by the aggregation of abnormal proteins that have classically been assessed post-mortem. The advent of *in vivo* imaging has led to the identification of various disease-relevant brain changes that can now be distinguished in living patients. While some *in vivo* changes are considered conclusive disease hallmarks, the relevance of others to Alzheimer’s disease specifically remains somewhat contentious. One such imaging hallmark where controversy remains is the appearance of hyperintense regions within white matter on T2-weighted MRI (in particular on fluid-attenuated inversion recovery (FLAIR) MRI), known as white matter hyperintensities (WMH). WMH are commonly reported and are considered by some to be a core feature of Alzheimer’s disease (Lee et al., 2016); however, the means by which these lesions contribute to disease-specific changes remains a topic of debate. The somewhat poor understanding of their disease relevance may stem from the limited information available when assessing these lesions *in vivo*.

Pathologically, WMH are characterised by a heterogeneous histological profile, including myelin pallor, myelin loss, axonal loss, gliosis and white matter infarction (Braffman et al., 1988; Fazekas et al., 1993; Gouw et al., 2008; Young et al., 2008; Schmidt et al., 2011a). Most of these histological changes are thought to be ischaemic in origin (Pantoni et al., 1996; Pantoni and Garcia, 1997; Topakian et al., 2010), and consequently, WMH have been proposed as a proxy measure for small vessel disease, and a surrogate endpoint for cerebrovascular clinical trials (Schmidt et al., 2004). While it may be a useful clinical surrogate, using global measures of WMH as a marker for small vessel disease disregards information about known pathological heterogeneity. Moreover, not all WMH observed on MRI have microangiopathic origin (Fazekas et al., 1993; McAleese et al., 2017), and different lesions may have distinct clinical and pathological correlates. Indeed, one limitation when investigating WMH *in vivo* is their homogeneous appearance on FLAIR MRI, which is nonspecific in distinguishing between variable underlying pathological changes. In the context of Alzheimer’s disease, distinguishing between different types of WMH is particularly relevant, given that some lesions are believed to be more closely associated with the disease, whereas others are thought to be less deleterious, age-associated injuries (Brickman et al., 2015; McAleese et al., 2017).

In an attempt to distinguish between WMH despite their homogeneous hyperintensity on FLAIR, classification schemes are commonly adopted, differentiating these lesions based on their location, shape or size. Commonly, these classification schemes distinguish periventricular WMH from deep WMH, or distinguish confluent lesions from punctate lesions. However, classification schemes differ, and the same terms are often used to define WMH in a disparate manner (Kim et al., 2008). For instance, while many visual rating scales define periventricular WMH as those lesions that have continuity with the lateral ventricles (Fazekas et al., 1987; Coffey et al., 1990; de Leeuw et al., 2000), others classify periventricular lesions based on their distance from the ventricular surface (Wen and Sachdev, 2004; DeCarli et al., 2005), or their shape or size (Schmidt et al., 1992; Scheltens et al., 1993). Unsurprisingly, the clinical and pathological correlates of different classes of WMH appear variable, given the inconsistency among classification schemes (van Straaten et al., 2006; Kim et al., 2008). Moreover, the use of categorical distinction to differentiate WMH itself has been criticised as somewhat arbitrary, as such classifications may not necessarily correspond to meaningful pathological differences (DeCarli et al., 2005).

*In vivo* methods that are able to identify and measure microstructural heterogeneity of these lesions could thus be highly valuable, given that they would likely reflect pathological differences among lesion types, above and beyond the binary identification of WMH that is possible with FLAIR. To this end, one MRI approach that is able to probe tissue microstructure is diffusion MRI, or diffusion-weighted imaging (DWI). Signal intensity on diffusion MRI is sensitive to the microscopic diffusion of water, and can be used to study white matter fibre architecture *in vivo*. As such, it is potentially sensitive to microstructure within WMH. The ability to appropriately model white matter structures, however, depends upon the type of diffusion data acquired, and the methods used to model these diffusion data. While diffusion tensor imaging (DTI) (Basser et al., 1994) is commonly used to model white matter microstructure, and has been widely applied to investigate microstructural properties of WMH, there are well-known limitations to the DTI model that render it problematic when interpreting results, particularly when multiple fibre orientations are present (Le Bihan et al., 2006; Jones, 2010; Jones et al., 2013).

Constrained spherical deconvolution (CSD) is a method that enables modelling of white matter in the presence of multiple fibre orientations (Tournier et al., 2004, 2007), even when there are crossing fibre populations within a voxel, and thus offers a means to model complex white matter structures better than the DTI model. However, the ability of CSD to model white matter may be confounded in areas where there is partial voluming with other tissues, such as grey matter (GM) or cerebrospinal fluid (CSF). A recently introduced variant of the CSD method, called single-shell 3-tissue CSD (SS3T-CSD) (Dhollander and Connelly, 2016a; Dhollander et al., 2016), is able to additionally estimate the GM and CSF compartments, minimising the effects of partial volume to more appropriately model white matter. More recently, SS3T-CSD has additionally been proposed as a means to provide insight into microstructural properties of pathological tissue (Dhollander et al., 2017). By characterising the diffusion signal obtained from tissue in terms of its relative composition of diffusion signal characteristics similar to those of the three distinct tissue types (i.e., those obtained from white matter (WM), GM and CSF), this method can provide insight into the microstructural properties of different types of tissue. This could then be applied to probe the underlying diffusional characteristics of WMH, and their potential microstructural heterogeneity *in vivo*.

In this study, we thus sought to investigate WMH using SS3T-CSD in a cohort of Alzheimer’s disease patients (*n* = 48) and healthy elderly control subjects (*n* = 94). Our aims were to investigate heterogeneity in the microstructural properties of these WMH *in vivo*, and to determine whether WMH exhibit distinct diffusional characteristics that would provide additional information to that obtained using conventional binary lesion classification schemes.

## 2 Materials and Methods

### 2.1 Participants

Participants included in this study were recruited as part of the Australian Imaging, Biomarkers and Lifestyle (AIBL) study of aging, and consisted of patients with clinical Alzheimer’s disease, and healthy elderly control subjects. Participants were classified into clinical groups according to AIBL criteria, and satisfied inclusion and exclusion criteria, as has been previously described (Ellis et al., 2009). Participants were included in this study if their MRI protocol included high b-value diffusion MRI, acquired at the Florey Institute of Neuroscience and Mental Health in Melbourne (*n* = 149). All participants also underwent an amyloid-β PET scan with ^11^C-PIB (carbon-11-labelled Pittsburgh compound B), and were classified as amyloid-positive or - negative based on a mean standardised uptake value ratio (SUVR) cut-off value of 1.4, as previously described (Rowe et al., 2013). Participants who had a clinical diagnosis of Alzheimer’s disease, but were amyloid-negative were excluded (*n* = 3). Participants were also excluded if they had incomplete demographic information (*n* = 2). FLAIR images were screened automatically for the presence of WMH. Subjects with substantial motion or intensity inhomogeneity artefacts on their FLAIR image were excluded from analysis, as automated segmentation of WMH on these subjects failed (*n* = 2). The final cohort included 142 subjects: 48 Alzheimer’s disease and 94 healthy control subjects. All subjects provided informed written consent, and the study was approved by the ethics committee at Austin Health.

### 2.2 Image Acquisition

MRI data were acquired for all subjects using a 3T Siemens Tim Trio System (Erlangen, Germany), with a 12-channel head coil receiver. DWI data were collected using echo planar imaging (EPI) with the following parameters: TR/TE = 9200/112 ms, 2.3 mm^3^ isotropic voxels, 128 × 128 acquisition matrix, acceleration factor = 2, diffusion-weighted images for 60 different gradient directions (b = 3000 s/mm^2^) and 5 volumes without diffusion-weighting (b = 0 s/mm^2^). FLAIR images were collected with the following parameters: 176 axial slices, voxel size 0.9 × 0.98 × 0.98 mm^3^, repetition time/echo time = 6000/420 ms, inversion time = 2100 ms, flip angle = 120°). A 3D MPRAGE (magnetization prepared rapid acquisition gradient echo) image (voxel size 1.2 × 1 × 1 mm^3^, repetition time/echo time = 2300/2.98, flip angle = 9°) was also obtained for each subject, and used to compute intracranial volume using SPM8 software. FLAIR and DWI data were then preprocessed and analysed using MRtrix3 (www.mrtrix.org) as described in the following sections, and as summarised in Figure 1.

**Figure 1:**
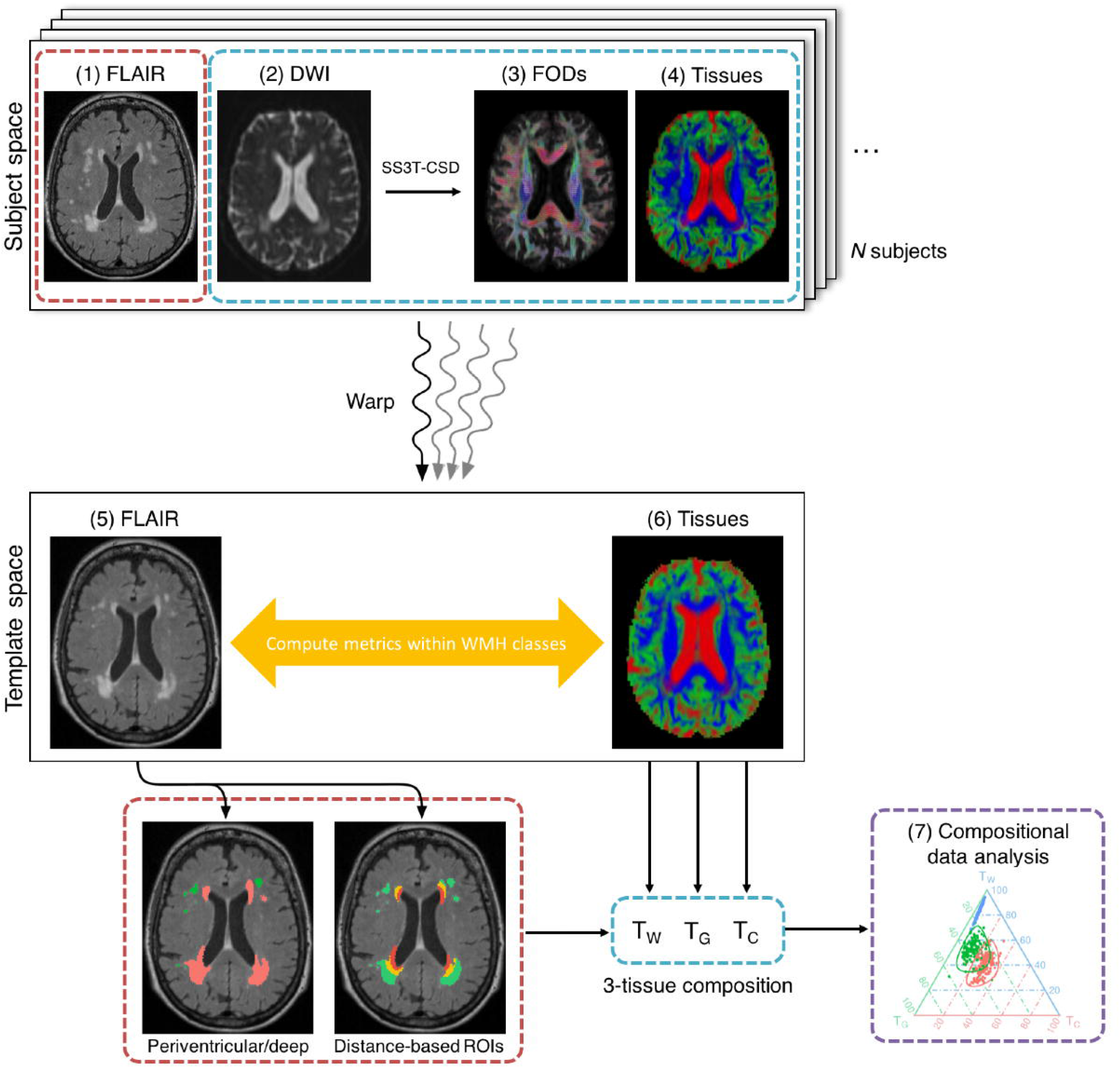
Schematic figure showing the major steps involved in diffusional analysis of white matter hyperintensities. (1) FLAIR and (2) DWI data were obtained for each subject, and were motion corrected and preprocessed. Single-shell 3-tissue constrained spherical deconvolution (SS3T-CSD) was performed on the DWI data to obtain (3) fibre orientation distribution (FOD) functions for white matter, as well as for the grey matter and CSF compartments, enabling (4) tissue maps to be created. Each subject’s images were warped to a common template space. Within each of the WMH classes obtained from classification schemes applied on (5) the FLAIR WMH segmentations, we then computed (6) the signal fractions obtained from the diffusion data (T_W_, T_G_, T_C_). The 3-tissue profile for each of the WMH classes could then be analysed with (7) compositional data analysis (CoDA).

### 2.3 Image Preprocessing

Preprocessing of diffusion-weighted images included denoising of data (Veraart et al., 2016), eddy-current distortion correction and motion correction (Andersson and Sotiropoulos, 2016), bias field correction (Tustison et al., 2010), and up-sampling resulting in 1.15×1.15×1.15mm^3^ isotropic voxels (Raffelt et al., 2012b). Intensity normalization across subjects was performed by deriving scale factors from the median intensity in select voxels of white matter, grey matter, and CSF in b = 0 s/mm^2^ images, then applying these across each subject image. Following these initial preprocessing steps, WM fibre orientation distribution (FOD) functions as well as GM and CSF compartments were computed using Single-Shell, 3-Tissue Constrained Spherical Deconvolution (SS3T-CSD), using group averaged response functions for WM, GM, and CSF obtained from the data themselves using an unsupervised method (Dhollander and Connelly, 2016a; Dhollander et al., 2016).

FLAIR images and MPRAGE images were also bias field-corrected. EPI susceptibility distortion correction of diffusion-weighted images was performed in conjunction with MPRAGE motion correction, using a registration-based method guided by a pseudo T1-contrast, which was estimated from the SS3T-CSD result (ie. the 3-tissue compartments) (Dhollander and Connelly, 2016b).

Spatial correspondence across subjects was achieved by first computing a group-specific population template via an iterative registration and averaging approach (Raffelt et al., 2011) using the white matter FOD images from 30 subjects from the study cohort. Each subject’s FOD image was then registered to the template via a FOD-guided non-linear registration (Raffelt et al., 2011, 2012a). FLAIR images were also corrected for motion (via registration to each subject’s MPRAGE image using ANTS (Avants et al., 2014)) and warped to the population template space, along with the WMH segmentations that were obtained from these FLAIR images (see next section).

### 2.4 Lesion Segmentation and Classification

WMH segmentations were performed automatically using the HyperIntensity Segmentation Tool (HIST) (Manjón et al., 2018). This automated tool segments WMH from 3D FLAIR images based on an ensemble of neural network classifiers. Given the variability of definitions used for periventricular versus deep classification of WMH in the literature, segmented WMH were then classified according to 2 different sets of criteria.

Firstly, WMH were classified either as “periventricular” or “deep” lesions, based on their distance and confluence with the lateral ventricles. WMH were classified as periventricular if the minimum distance from the lateral ventricles was less than 5.0 mm; or, if the average distance of a continuous lesion was less than 20.0 mm. WMH were otherwise classified as deep.

Secondly, we performed a classification of each WMH voxel separately based on its distance from the ventricles, regardless of continuity of a lesion (see Fig. 2), extending the distance-based classification scheme described by DeCarli et al. (2005). We defined three regions-of-interest by dilating a ventricular mask obtained from the population template image to an area that included anything 10 mm or less from the lateral ventricles in all directions. WMH voxels falling within this region were classified as Region 1, regardless of whether they had contiguous WMH voxels that extended beyond this region. A second region (Region 2) was defined by further dilating the first region to obtain an area between 10 mm and 20 mm from the lateral ventricles. The third region (Region 3) consisted of all remaining white matter beyond 20 mm from the lateral ventricles.

**Figure 2:**
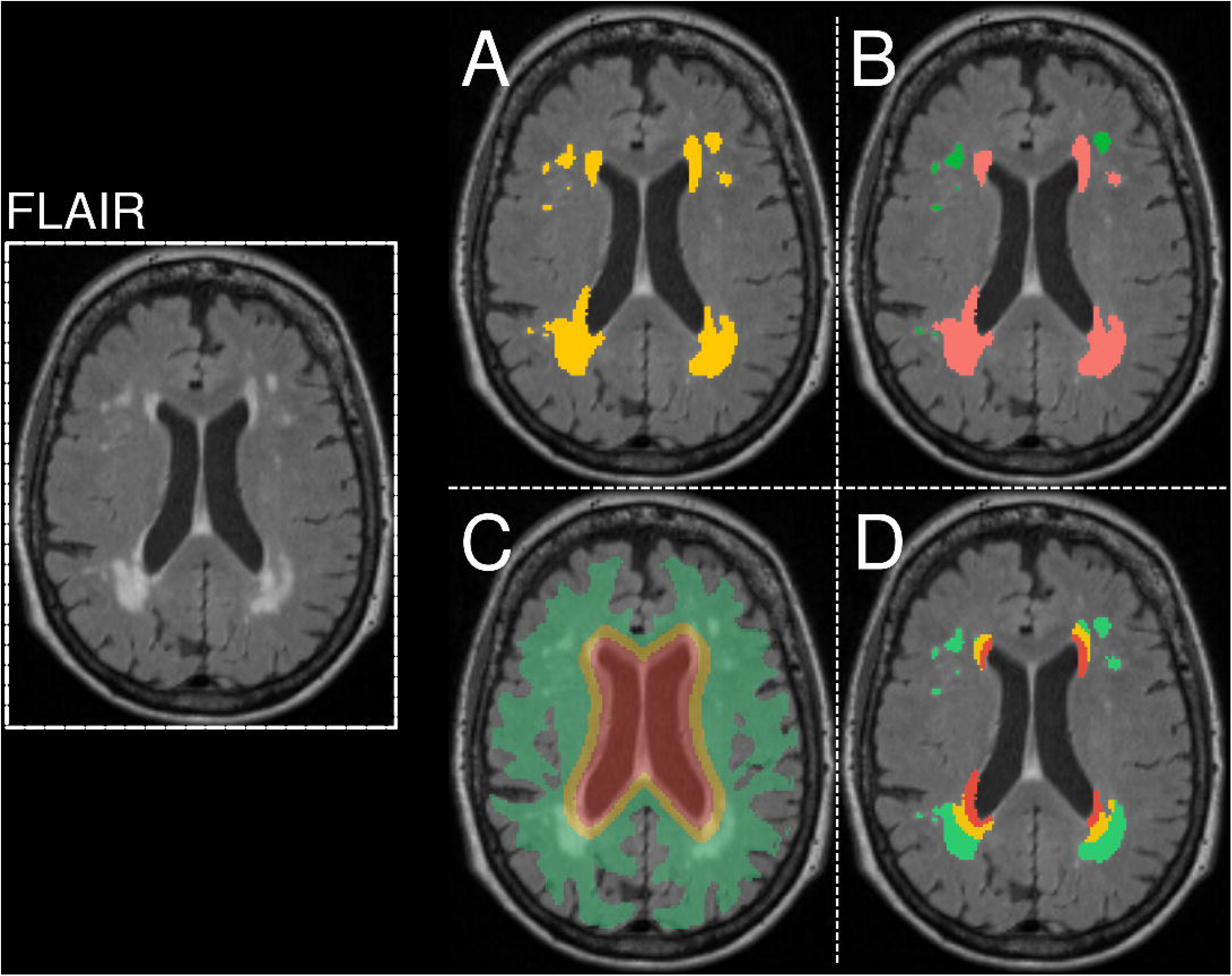
White matter hyperintensity segmentation and classification schemes. White matter hyperintensities were automatically segmented from FLAIR MRI. A single axial slice of the FLAIR image is shown on the left for a given subject in template space. (A) The WMH segmentation for this subject is shown in yellow. (B) An example of the periventricular/deep classification scheme is shown for the WMH in a given subject. WMH were classified as “periventricular” if the minimum distance from the lateral ventricles was less than 5.0 mm in subject space or if the average distance of a continuous lesion was less than 20.0 mm. WMH were otherwise classified as “deep”. (C) Regions of interest were defined by repeatedly dilating a ventricular mask obtained from the population template brain. The first region (Region 1) is shown in red, and included an area 10mm or less from the lateral ventricles in all directions. Region 2 is shown in yellow, and was defined by further dilating Region 1 to obtain an area between 10mm and 20mm from the lateral ventricles. Region 3, shown in green, consisted of all remaining white matter beyond 20mm from the lateral ventricles. (D) Lesions were classified into these distance based classes according the Region with which they overlapped.

In addition to the above classification schemes, WMH voxels were also classed based on the brain lobe in which they were located, as described in the Supplementary Material.

We computed a normal-appearing white matter (NAWM) mask for each subject in template space. A WM segmentation was obtained from each subject’s T1 image in template space, using FSL FAST (Zhang et al., 2001). The WMH for each subject were subtracted from that subject’s WM segmentation, and the remaining NAWM mask was subsequently eroded by one voxel in three dimensions to ensure that voxels within this mask represented normal-appearing WM only.

### 2.5 Representing the composition of WMH and NAWM as 3-tissue diffusion signal fractions

The WM, GM and CSF compartments were obtained from SS3T-CSD as described above. As CSD techniques typically operate directly on the *absolute* diffusion-weighted signal, these compartments are also directly proportional to the *absolute* amount of diffusion-weighted signal attributable to each of these tissue types in each voxel. In previous work, it was found that certain microstructural aspects of WM pathology result in diffusion-weighted signals akin to those represented by the response functions measured from GM and CSF (Dhollander et al., 2017). In this context, these were referred to as “GM-like” and “CSF-like” (diffusion-weighted) signals. In this work, we intended to study the *relative* makeup of the diffusion-weighted signal in terms of WM-, GM- and CSF-like tissue signal *fractions*. To distinguish these from the *absolute* signals, we refer to the 3-tissue signal *fractions* as T_W_, T_G_ and T_C_ respectively.

To obtain T_W_, T_G_ and T_C_ for each WMH category and NAWM per subject, we first computed the absolute WM-like, GM-like and CSF-like signal over each WMH category, and normalised the resulting triplet of absolute signals to sum to one. The resulting 3-tissue composition of each WMH category and NAWM for each subject included in the analysis was plotted on a ternary plot. Ternary plots were created using the *ggtern* package in R (Hamilton, 2018), enabling easy visualisation of the 3-tissue profile of the WMH categories and NAWM regions.

### 2.6 Statistical analysis

Demographic variables and WMH volumes were compared between the two clinical groups. We used *t*-tests and *χ*^2^ tests to compare age, sex, and intracranial volume. WMH volumes were compared between groups by performing ANCOVAs, including age and intracranial volume as covariates. Given that the distribution of WMH volumes across subjects is right-skewed, we performed a cubic root transformation to all WMH volumes to normalise the distribution. Bonferroni-corrected *P*-values were used to determine statistical significance.

We compared the 3-tissue profiles between different classes of WMH and NAWM, across all subjects. However, statistical analysis on such 3-tissue compositions is not trivial: boundedness (0 < T_W_ < 1; 0 < T_G_ < 1; 0 < T_C_ < 1) and non-independence (T_W_ + T_G_ + T_C_= 1) render traditional statistical analysis of the 3-tissue compositions inappropriate. Therefore, we adopted the Compositional Data Analysis (CoDA) framework (Aitchison, 1982; Pawlowsky-Glahn and Buccianti, 2011). To this end, we designed an isometric log-ratio transform (Egozcue et al., 2003) tailored to our 3-tissue compositions at hand:

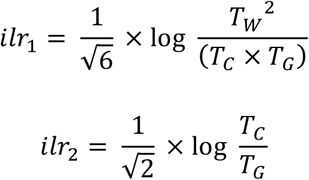

Unlike the original 3-tissue compositions themselves, which are bounded and non-independent, the isometric log-ratio transformed data are free to range across all real numbers and are independent, resulting in only 2 degrees of freedom in the 3-tissue compositional space. Classical multivariate methods can then be applied to the log-ratio transformed data to perform statistical analysis (Martin-Fernández et al., 2015).

We performed MANOVAs on the isometric log-ratio transformed data to determine whether different classes of WMH were statistically significantly different from one another. A MANOVA was performed between each pair of classes separately for all three analyses, and Bonferroni corrections were applied within each analysis to correct for the multiple comparisons performed. Pillai’s trace was used as the multivariate test in all analyses.

Bivariate plots of the isometric log-ratio transformed data, which exhibited the confidence ellipses and ellipse centre of each WMH class and NAWM, were created using the *ggplot2* package in R (Fox and Weisberg, 2011).

### 2.7 Data and code availability statement

The data on which the findings of this study are based were collected as part of the AIBL study, for which all participants gave written and informed consent to participate for the purposes of the study. As such, raw data remain confidential. However, data may be made available upon reasonable request from AIBL (https://aibl.csiro.au/research/support). The code required for the SS3T-CSD analysis is currently being prepared for release within MRtrix (www.mrtrix.org), and will be publicly available as soon as the process is complete.

## 3 Results

### 3.1 Clinical and demographic characteristics

Clinical and demographic characteristics for the Alzheimer’s disease and healthy control groups are summarised in Table 1. No significant differences were observed between groups for age, sex, or intracranial volume.

**Table 1:** Descriptive statistics

|  | **Alzheimer's disease (n = 48)** | **Healthy elderly controls (n = 94)** | **Statistics** | **p-value** |
| --- | --- | --- | --- | --- |
| **Age, mean (SD)** | 77.42 (8.3) | 78.12 (7.4) | $t(140) = 0.75$ | 0.46 |
| **Males (%)** | 21 (43.8) | 44 (46.8) | $\chi^2(1) = 0.03$ | 0.87 |
| **11C-PIB positivity (%)** | 48 (100) | 31 (33) | $\chi^2(1) = 57.83$ | < 0.001 |
| **Intracranial volume (cm<sup>3</sup>), mean (SD)** | 1402.95 (128.1) | 1433.61 (134.8) | $t(140) = 1.30$ | 0.20 |
| **Total WMH volume (cm<sup>3</sup>), mean (SD)** | 13.90 (13.3) | 8.43 (9.4) | $t(71.4) = 2.54$ | 0.01 |
| **Regional WMH volumes (cm<sup>3</sup>), mean (SD)** |  |  |  |  |
| **Periventricular** | 12.81 (12.7) | 7.54 (8.8) |  |  |
| **Deep** | 1.09 (1.25) | 0.89 (1.2) |  |  |
| **Region 1** | 2.61 (2.1) | 2.32 (2.0) |  |  |
| **Region 2** | 4.49 (3.4) | 2.50 (2.6) |  |  |
| **Region 3** | 6.72 (8.7) | 3.57 (5.6) |  |  |
| **Cubic root WMH volumes, mean (SD)** |  |  |  |  |
| **Total WMH** | 2.18 (0.73) | 1.83 (0.63) | $F(1,138) = 13.75$ | < 0.001 |
| **Periventricular WMH** | 2.11 (0.74) | 1.74 (0.64) | $F(1, 138) = 14.69$ | < 0.001 |
| **Deep WMH** | 0.91 (0.35) | 0.83 (0.35) | $F(1, 138) = 1.38$ | 0.24 |
| **Region 1 WMH** | 1.25 (0.42) | 1.22 (0.38) | $F(1, 138) = 0.48$ | 0.49 |
| **Region 2 WMH** | 1.51 (0.50) | 1.19 (0.47) | $F(1, 138) = 24.06$ | < 0.001 |
| **Region 3 WMH** | 1.62 (0.69) | 1.26 (0.61) | $F(1,138) = 14.81$ | < 0.001 |

### 3.2 White matter hyperintensity volume

Total WMH volume was higher in Alzheimer’s disease patients compared to control subjects, after controlling for age and intracranial volume (*F*(1,138) = 13.75, *P* < 0.001). Considering our two WMH classification schemes in turn: (a) No significant differences were observed between Alzheimer’s disease and control groups for deep WMH volume (*F*(1,138) = 1.38, *P* = 0.24), while there was a significantly greater periventricular WMH volume in Alzheimer’s disease patients compared to controls (*F*(1,138) = 14.69, *P* < 0.001). (b) Alzheimer’s disease patients had a comparable load of WMH within Region 1 when compared to healthy elderly control subjects (*F*(1,138) = 0.48, *P* = 0.49). However, the WMH load in Region 2 and *Region 3* was significantly higher in patients compared to controls (*Region 2 F*(1,138) = 24.06, *P* < 0.001; *Region 3*: *F*(1,138) = 14.81, *P* < 0.001). Each of the above was compared between amyloid-positive and amyloid-negative healthy control subjects, but no significant differences were found in WMH volumes for any of these classes (see Supplementary Table 1).

Differences in the lobar volumes between patients and controls are described in the Supplementary Material, along with the 3-tissue compositional analysis for lobar WMH.

### 3.3 Comparing 3-tissue profiles of WMH classes

#### 3.3.1 Periventricular vs Deep WMH

As shown in the boxplots in Figure 3, and ternary plot in Figure 4, the 3-tissue profiles derived from SS3T-CSD showed different compositions of T_W_, T_G_, and T_C_ in periventricular and deep WMH, with higher T_C_ in periventricular WMH than in deep WMH. Periventricular and deep WMH formed distinct clusters based on their 3-tissue profiles alone, as did NAWM. The log-ratio transformed data is shown in Figure 5, which exhibits the mean and 95% confidence ellipses of each group (WMH class or NAWM) in the 2-dimensional coordinate system, revealing the distinct clusters formed by each group. Statistical analysis using a MANOVA showed a statistically significant difference between NAWM and periventricular WMH in terms of 3-tissue composition (*F*(2, 281) = 5336.79, *P* < 0.001, Pillai’s trace = 0.974), between NAWM and deep WMH (*F*(2, 278) = 1029.65, *P* < 0.001, Pillai’s trace = 0.881), and between periventricular and deep WMH (*F*(2, 278) = 275.58, *P* < 0.001, Pillai’s trace = 0.665), with a Bonferroni corrected significant *P*-value of 0.0167.

**Figure 3:**
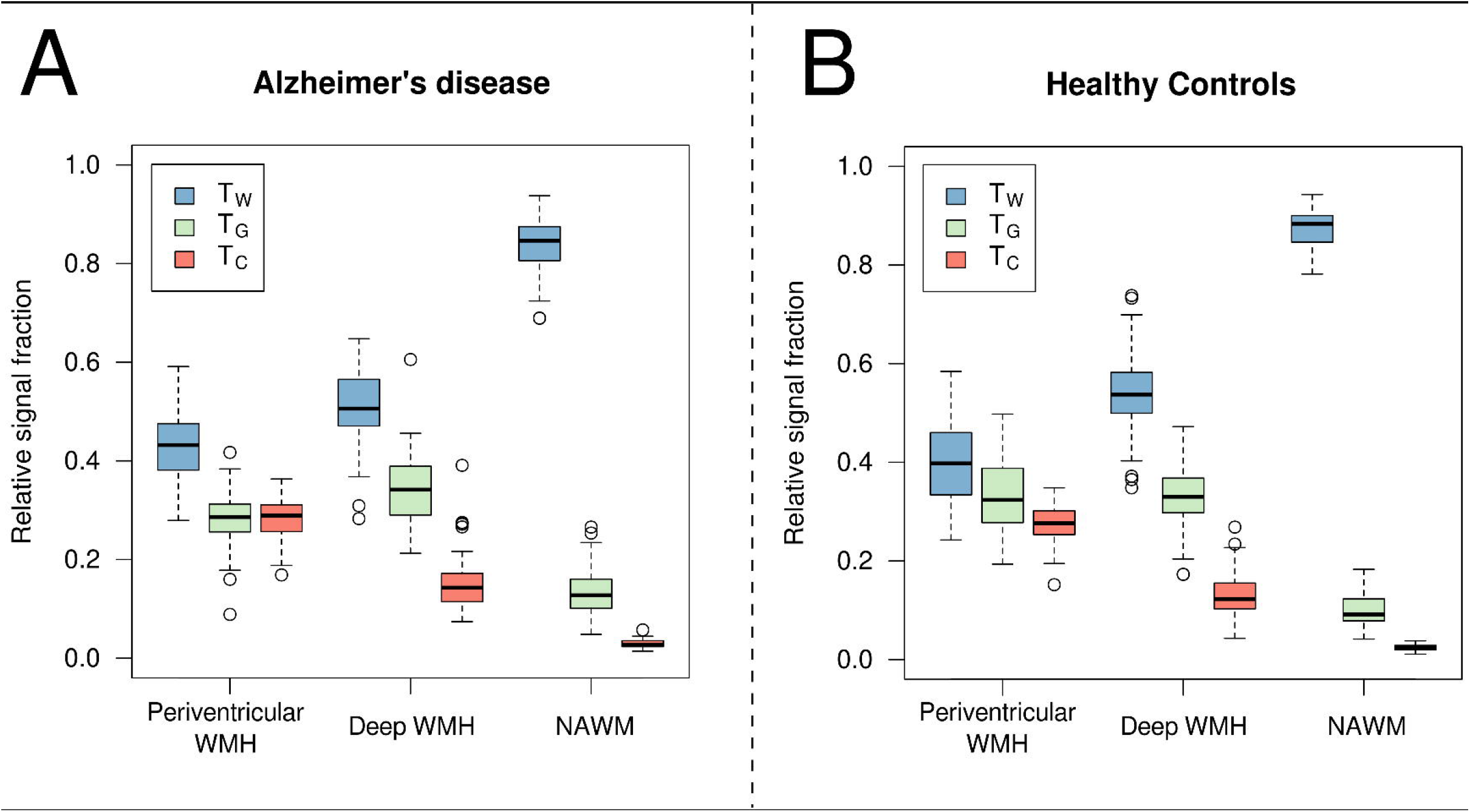
Boxplots showing relative signal fractions within lesions and NAWM. The relative T_W_, T_G_, T_C_ signal fractions are displayed across all (A) Alzheimer’s disease subjects (*n* = 48) and (B) healthy elderly control subjects (*n* = 94) as the median, first, and third quartiles, and 95% confidence interval of the median. Normal-appearing white matter (NAWM) exhibits high T_W_ fraction as expected, reflecting the high white matter-like diffusion profile with relatively low T_G_ and T_C_ fractions, both in healthy elderly and Alzheimer’s disease subjects. In contrast, the WMH classes exhibit higher T_G_ and T_C_ fractions. Periventricular and deep WMH can be distinguished from one another by their relative signal fractions, and the two lesion types exhibit a similar profile in Alzheimer’s disease patients as they do in control subjects.

**Figure 4:**
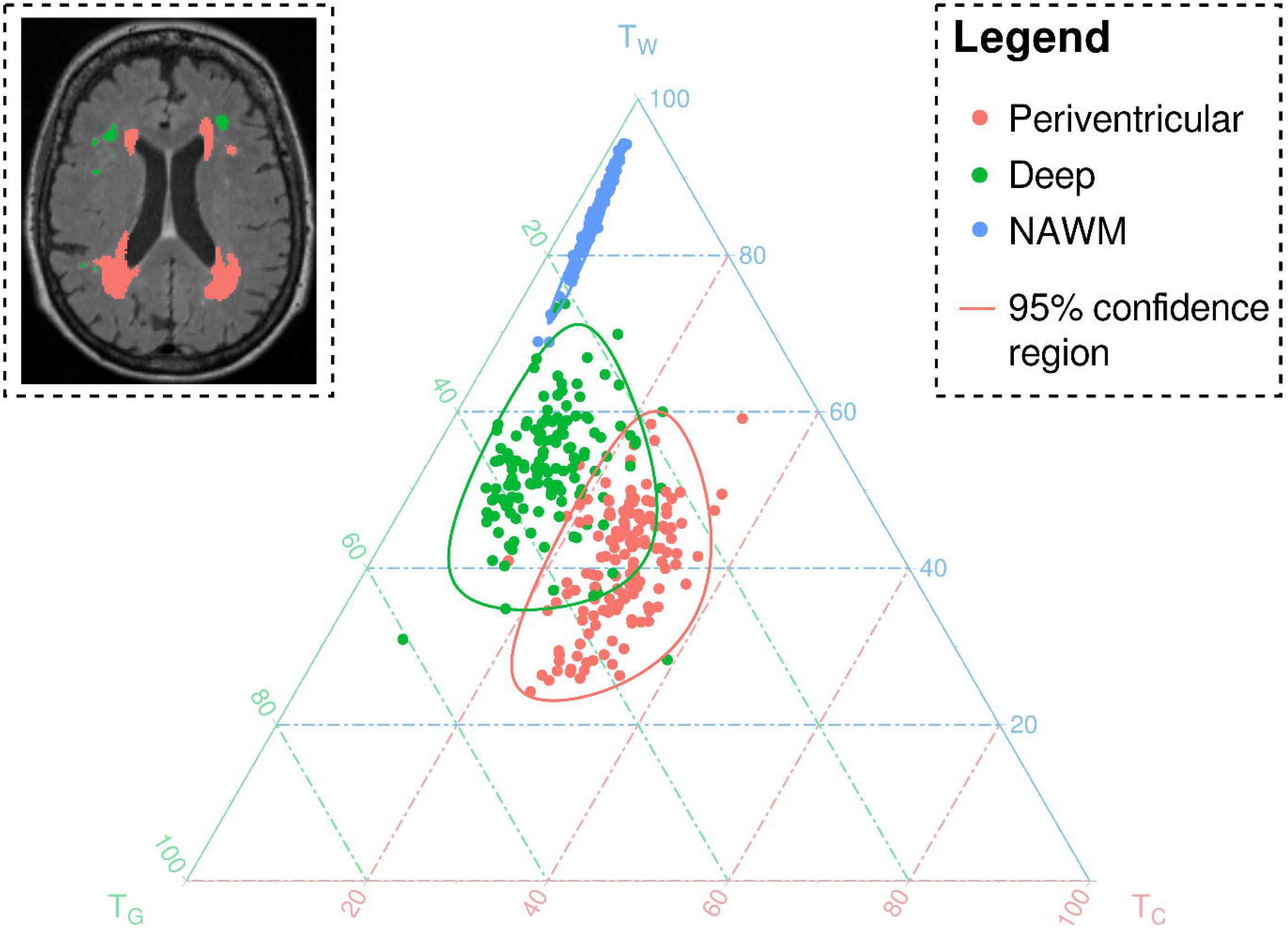
Ternary plot exhibiting relative signal fractions within periventricular and deep lesions and NAWM. For each subject, the periventricular WMH, deep WMH, and NAWM are displayed on a ternary plot, with the location of the data point corresponding to the relative T_W_, T_G_ and T_C_ fractions of the lesions (or NAWM) for that subject. Given the similarity in the profile of lesions in Alzheimer’s disease subjects and controls, all subjects are included here. The relative tissue fraction is shown as a percentage along the left (T_W_), right (T_G_), and bottom (T_C_) axes. Remarkably, the periventricular WMH, deep WMH, and NAWM appear in distinct clusters, exhibiting their different profiles with regard to relative tissue fractions obtained from the SS3T-CSD diffusion data. An example of the classification of lesions into periventricular and deep is shown in the inset. The 95% population confidence regions are plotted for each WMH class and NAWM.

**Figure 5:**
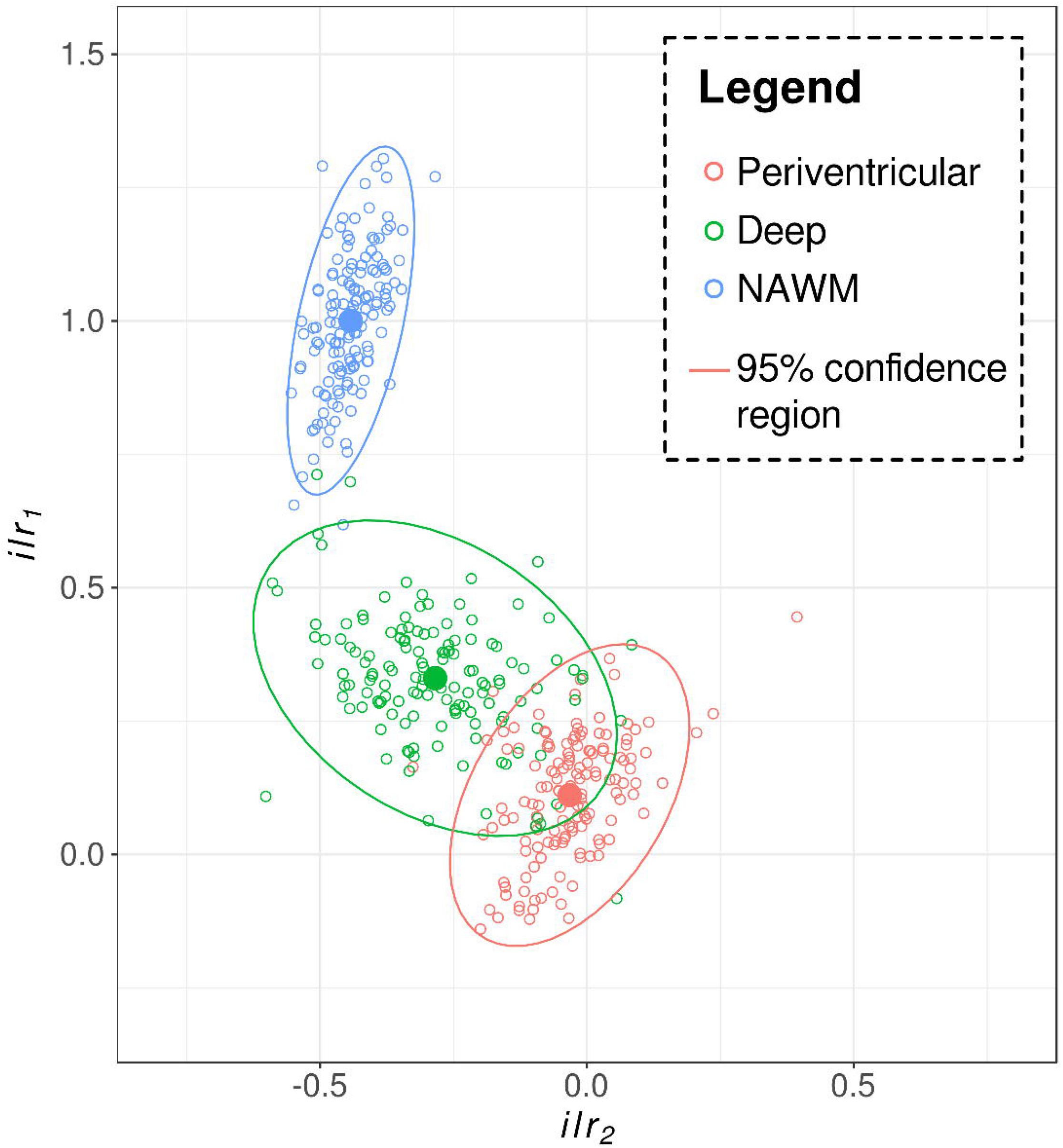
Isometric log-ratio transformed diffusional data comparing periventricular WMH, deep WMH and NAWM. An isometric log-ratio (*ilr*) transform was applied to the relative signal fractions (T_W_, T_G_, T_C_), to transform the compositional data into a two-coordinate system. The transformed data for the periventricular and deep WMH, and NAWM are shown here, and reflect the same data shown in Figure 4. The centroid for each WMH class or NAWM is shown as a solid circle, while the solid line reflects the 95% population confidence ellipse for each class. Statistical analyses comparing the diffusional profile of lesion classes and NAWM was performed on this isometric log-ratio transformed data. Pairwise MANOVAs exhibited that there were statistically significant differences in the transformed data between the two lesion types, and between the lesions and NAWM. This could then be meaningfully back-transformed to interpret a significant difference in the mean diffusion profile between the two lesion types, and between the lesions and NAWM.

#### 3.3.2 Distance-based region analysis

Figure 6A shows a ternary plot exhibiting the 3-tissue profiles of WMH within three distance-based regions, defined by concentric distances from the lateral ventricles. As shown in the ternary plot, the different region classes of WMH again exhibited distinct 3-tissue profiles, and differed from NAWM. The log-ratio transformed data is shown in Figure 6B. Multivariate tests using a MANOVA showed a significant difference between NAWM and all three region classes (Region 1: *F*(2, 281) = 5416.33, *P* < 0.001, Pillai’s trace = 0.975; Region 2: *F*(2, 281) = 1706.83, *P* < 0.001, Pillai’s trace = 0.924; Region 3: *F*(2, 281) = 972.31, *P* < 0.001, Pillai’s trace = 0.874), as well as significant pairwise differences between each of the region classes themselves (Region 1 vs Region 2: *F*(2, 281) = 485.80, *P* < 0.001, Pillai’s trace = 0.776; Region 1 vs Region 3: *F*(2, 281) = 740.50, Pillai’s trace = 0.841; Region 2 vs Region 3: *F*(2, 281) = 52.28, *P* < 0.001, Pillai’s trace = 0.271), with a Bonferroni-corrected significant *P*-value of 0.0083.

**Figure 6:**
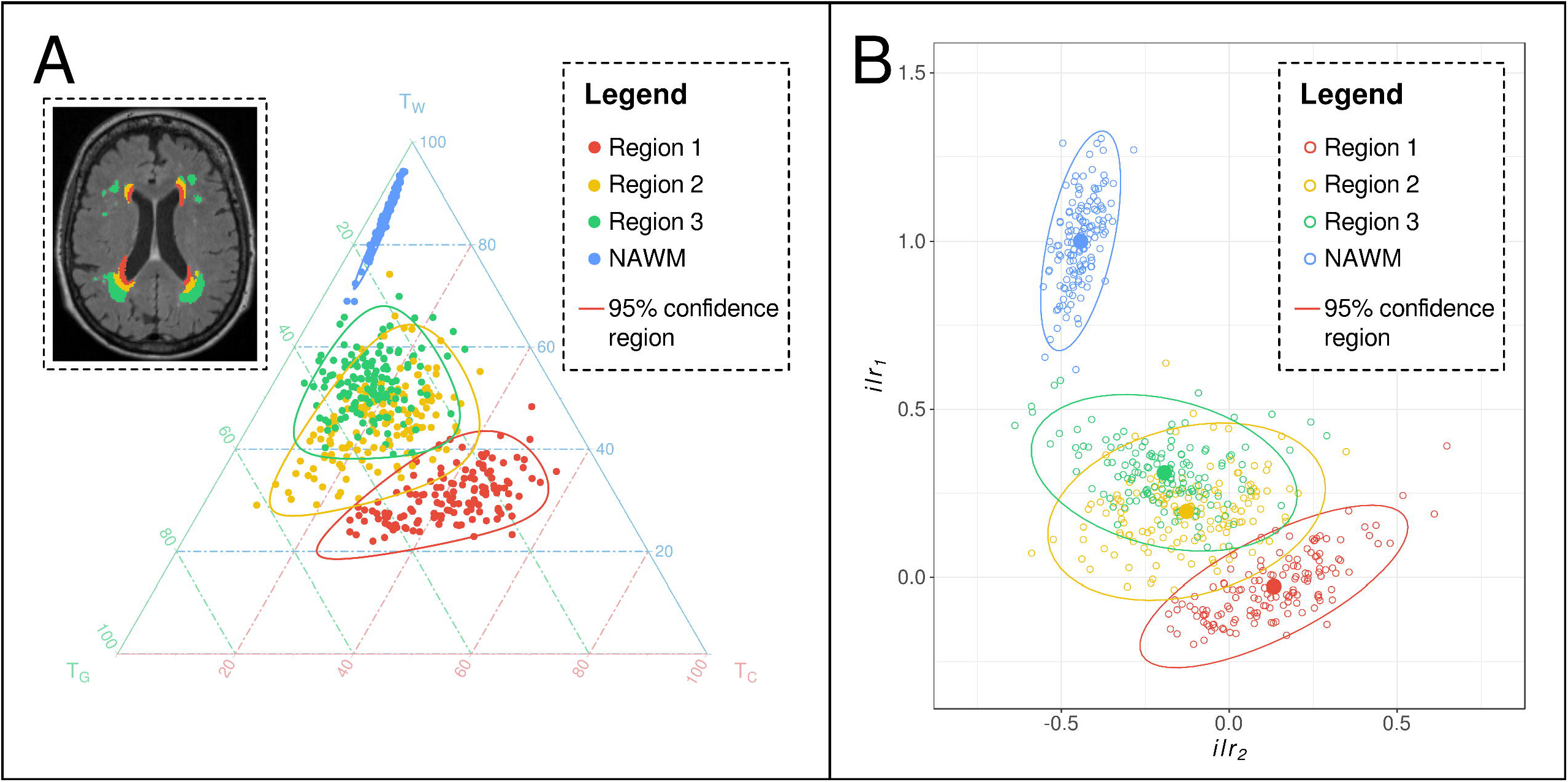
Compositional data analysis of lesions in each distance-based region and NAWM. (A) As with Figure 4, the relative T_W_, T_G_ and T_C_ fraction for WMH or NAWM is displayed on a ternary plot. Each data point reflects the mean T_W_, T_G_ and T_C_ fraction across all voxels falling within the NAWM mask (blue), or within one of three regions-of-interest (Regions 1, 2, and 3), located at concentric distances from the lateral ventricles, from a single subject. When looking across all subjects, the WMH falling within each region-of-interest exhibit a distinct diffusional profile, and the diffusional profile of each lesion area can be distinguished. The lines reflect the 95% confidence region or predictive region for each lesion region or NAWM. (B) An isometric log-ratio transform was applied to the relative signal fractions (T_W_, T_G_, T_C_) to transform the compositional data into a two-coordinate system. Each data point reflects the transformed diffusion profile from a single subject, while the solid circle reflects the centroid and concentric solid line reflects the 95% population confidence ellipse for each lesion region or NAWM. Statistical analysis was performed on the isometric log-ratio transformed data, which showed significant pairwise differences in the diffusional profile between all regions, and between NAWM and each region.

## 4 Discussion

In this study, we applied novel diffusion MRI methods to investigate *in vivo* microstructural characteristics of white matter hyperintensities in a cohort of healthy elderly and Alzheimer’s disease subjects. The findings of this study are of twofold significance: firstly, they demonstrate that different classes of WMH can be differentiated from one another based on their microstructural properties; and secondly, they demonstrate the ability of this novel diffusion MRI method to probe *in vivo* heterogeneity within WMH that would otherwise appear homogeneous. These findings suggest that the 3-tissue profiles utilised in this study could be used to probe WMH *in vivo* as heterogeneous entities, which could enable further understanding of their clinical and pathological association with Alzheimer’s disease.

### 4.1 Periventricular and deep WMH characterised by distinct 3-tissue profiles

WMH are commonly dichotomised into periventricular and deep lesion subtypes, and although this categorical distinction is argued to be somewhat arbitrary (DeCarli et al., 2005), these two lesion classes have been suggested to have differing neuropathological substrates (Fazekas et al., 1991, 1993). As such, we hypothesised that periventricular and deep lesions would exhibit different microstructural properties, and that these differences could be detected with diffusion MRI, despite appearing visually homogeneous on FLAIR. Utilising an automated classification method that differentiated confluent periventricular from deep WMH, our data indicate that these two lesion classes exhibit distinct 3-tissue profiles in our cohort of healthy elderly and Alzheimer’s disease subjects. As shown in Figure 4, the relative T_W_-T_G_,-T_C_ compositions of periventricular and deep WMH across subjects could be clearly distinguished from one another.

Alzheimer’s disease patients in our cohort exhibited a significantly higher periventricular WMH volume when compared to healthy elderly subjects. This is consistent with the increased severity of extensive periventricular hyperintensities that has been previously reported in Alzheimer’s disease (Barber et al., 1999). Moreover, periventricular WMH, rather than deep WMH, have been preferentially associated with cognitive impairment and dementia (O’Brien et al., 1996; de Groot et al., 2002; Prins et al., 2004). As such, these lesions could be more deleterious than deep WMH, and more closely associated with Alzheimer’s disease symptomatology (Fazekas et al., 1987). The 3-tissue profile of these lesions could thus itself be a useful reflection of more adverse underlying pathology, and could potentially provide an *in vivo* probe to distinguish more harmful, potentially disease-related changes, from benign age-related processes, regardless of the size, shape or location of a lesion.

Histologically, confluent periventricular lesions are characterised by substantial axonal and myelin loss and reactive gliosis, whereas punctate deep WMH have been reported to exhibit more mild changes with myelin loss (Fazekas et al., 1991, 1993; Schmidt et al., 2011a). In our work, the periventricular WMH exhibited distinctively higher T_C_ than NAWM and deep WMH. This suggests an increase in free fluid, given that the model for T_C_ is derived from the diffusion signal in pure CSF voxels, and hence an increase in T_C_ reflects a shift towards more fluid-like properties. Such a finding is in line with histological work, as the substantial myelin and axonal loss that presumably arises within the periventricular lesions is likely accompanied by increased extracellular fluid (Weller, 1998). On the other hand, the 3-tissue profile of deep WMH across subjects suggests less severe damage: while it was clearly distinguishable from that of normal-appearing white matter, it was characterised by decreased T_W_ and relative increases both in T_G_ and T_C_, rather than a marked increase in T_C_ as was observed in periventricular WMH. This suggests that disruption to white matter within these deep WMH may reflect less severe changes, in line with the mild myelin loss and gliosis that is histologically observed.

Importantly, these two WMH types were distinctively different in 3-tissue composition when compared to normal-appearing white matter across both healthy elderly subjects and Alzheimer’s disease patients. NAWM across subjects had high T_W_ content as expected, reflecting a 3-tissue profile that was similar to healthy white matter. The observed distribution of NAWM profiles across subjects (Fig. 4) suggests that in some individuals, even non-lesional white matter was exhibiting signs of some pathological insult (though more subtle and likely widespread), which is as expected given our previous findings of substantial fibre tract degeneration in the Alzheimer’s disease patients from the same cohort (Mito et al., 2018). Indeed, NAWM surrounding WMH has been shown in other studies to exhibit diffusional abnormalities (Maillard et al., 2014; Maniega et al., 2015, 2018), contributing to a so-called WMH “penumbra” (Maillard et al., 2011).

### 4.2 3-tissue profile within lesion regions varies with distance from ventricles

It should be noted, however, that heterogeneity in the 3-tissue profile of lesions could be observed *within* confluent periventricular lesions (Fig. 7) and as such, averaging the tissue profile over the whole lesion class could potentially fail to reflect variability in the microstructural changes within these continuous lesions. For instance, lesions within the immediate periventricular zone likely have distinguishable characteristics from large confluent hyperintensities (Sze et al., 1986; Fazekas et al., 1993; Schmidt et al., 2011a), despite potentially becoming continuous with them over time. The increased T_C_ observed in the periventricular lesions could have been driven by the increased interstitial fluid resulting from ependymal discontinuation in the immediate periventricular zone, and subsequent CSF leakage into the white matter, rather than being a characteristic of the whole confluent lesion. We were thus interested in extending the above analysis that was based on a periventricular/deep classification with a complementary investigation that probed the 3-tissue profile of lesion areas based on their distance from the lateral ventricles.

**Figure 7:**
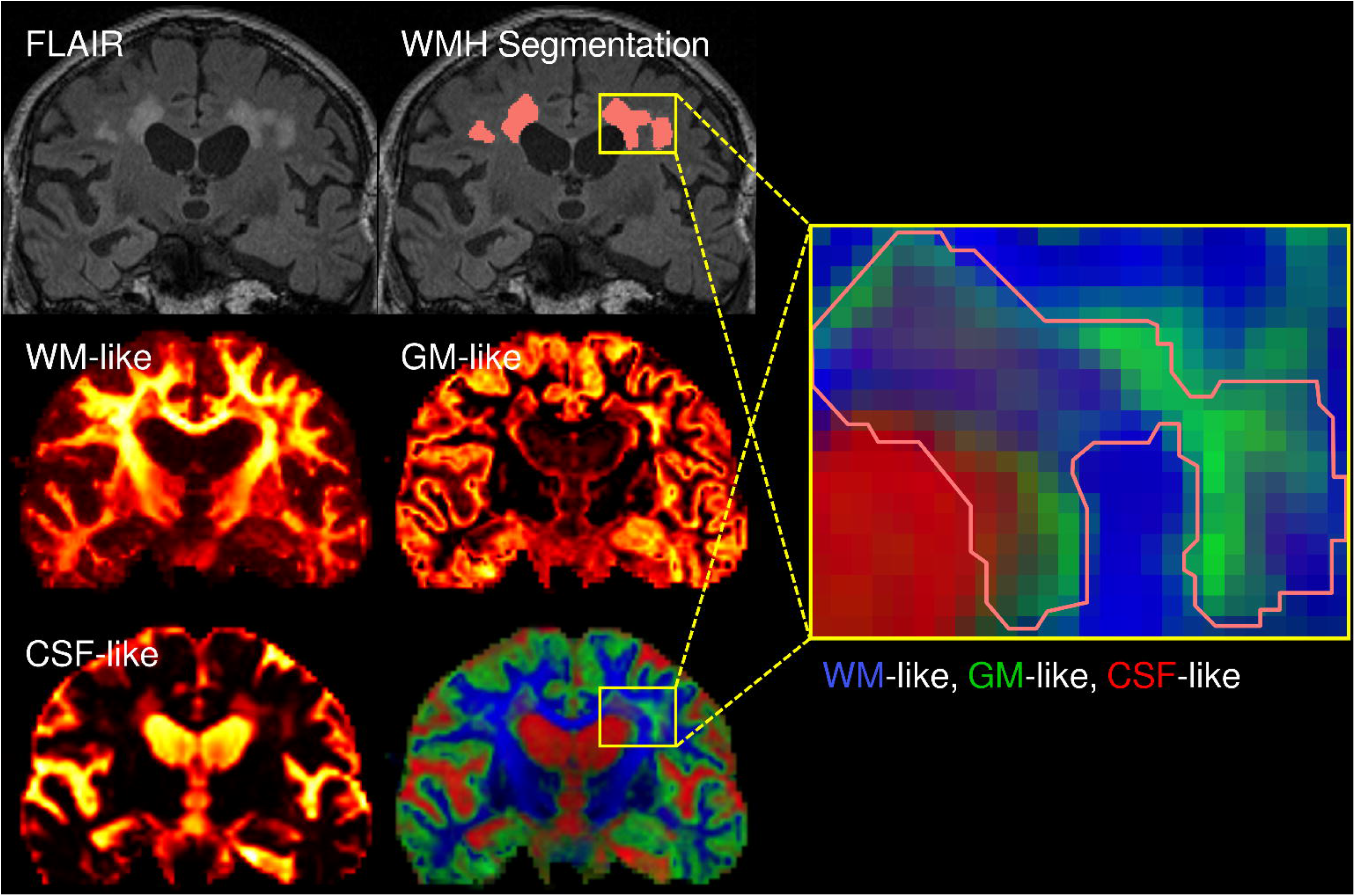
Investigating in vivo microstructural heterogeneity of white matter hyperintensities using diffusion MRI. WMH appear as homogeneous lesions on FLAIR MRI (top left), from which they can be segmented (segmentation shown on the FLAIR image on top right in pink). We computed the white matter-like (WM-like), grey matter-like (GM-like) and CSF-like signal compartments using single-shell, 3-tissue constrained spherical deconvolution (SS3T-CSD) (shown as heat maps in middle left, right, and bottom left, respectively). The three tissue compartments are shown in a single tissue-encoded colour map on the bottom right (WM = blue, GM = green, CSF = red). As can be seen in inset on the right, the region corresponding to WMH shows a heterogeneous mix of the three tissue compartments (segmentation outline shown in pink).

Here, we defined three regions-of-interest at set distances from the ventricles. While others have similarly used arbitrary distances to classify periventricular and deep WMH (Wen and Sachdev, 2004; DeCarli et al., 2005), we should highlight that our objective here was not to investigate different “classes” of lesions, but rather to determine if there were consistent, distance-related characteristics to WMH sub-regions that would potentially reflect heterogeneity within confluent lesions. Our findings suggest that lesion areas within each distance band from the lateral ventricles indeed exhibited distinct microstructural characteristics. The high T_C_ observed in lesion regions falling within 10 mm of the lateral ventricles (within Region 1) suggests that WMH areas within this immediate periventricular zone have increased free fluid content when compared to NAWM. This could arise due to increased interstitial fluid as a consequence of partial loss of the ependymal lining, which is characteristic of periventricular lesions in elderly individuals (Fazekas et al., 1993).

The overall lesion volume within this most proximal distance band was comparable between healthy elderly individuals and Alzheimer’s disease patients in our cohort, which suggests that lesions in this immediate periventricular area reflect benign, age associated changes that likely arise due to their location in a potential watershed region. This appears to contradict the suggestion that in periventricular lesions (using the periventricular/deep WMH classification above), a 3-tissue profile characterised by high T_C_ is indicative of deleterious underlying pathology due to there being a greater volume of periventricular lesions in Alzheimer’s disease subjects than in controls. In fact, when lesion regions were classified according to distance from the ventricles irrespective of whether they were contiguous with other lesion regions, it was those regions falling within Regions 2 and 3 (which had distinctively lower T_C_ than those lesion region falling within 10 mm of the ventricles) that were greater in volume in Alzheimer’s disease subjects than in controls.

These findings, though seemingly contradictory with the aforementioned findings using a periventricular/deep classification, suggest an important consequence: the use of conventional classification schemes (particularly distinguishing between periventricular and deep lesion types) could be misleading, as they do not take account of the substantial heterogeneity within white matter lesions. While the arbitrary nature of conventional classification schemes, and the importance of distinctions among different types of periventricular and deep WMH have previously been highlighted (DeCarli et al., 2005; Kim et al., 2008; Schmidt et al., 2011b), most research studies have adopted conventional classifications to probe disease-relevant associations.

Inferences are commonly made regarding the clinical relevance and pathological correlates of WMH based on these conventional classes, and could similarly be made from our own analysis above; however, careful consideration is required of the resultant findings given the aggregation of complex information into a single, oversimplified class. The use of three distance-based classes reveals that different conclusions can be drawn from the same data by investigating the lesions while considering the heterogeneity that may be present at different distance bands. Importantly, this highlights the pitfalls of using conventional classification schemes, particularly when investigating their relevance to disease. Indeed, we suggest that one should not consider how lesion types are associated to disease without considering their existing heterogeneity, which we propose can now be investigated *in vivo*.

### 4.3 Probing microstructure with diffusion MRI

In addition to identifying microstructural differences between WMH in different locations and classes, an important result of the present study was in establishing the feasibility of diffusion MRI, and particularly SS3T-CSD, in assessing within-lesion microstructural heterogeneity *in vivo*. As suggested above, the somewhat limited understanding of the contribution of WMH to Alzheimer’s disease could stem from the simplistic way in which these lesions are commonly investigated *in vivo*, despite post mortem evidence that they are pathologically complex. To this end, we suggest that diffusion MRI could function as an *in vivo* probe to assess WMH, potentially in conjunction with FLAIR MRI, to investigate how these lesions may contribute both clinically and pathologically to Alzheimer’s disease.

Previous diffusion MRI research has attempted to investigate microstructural changes within WMH, by investigating these lesions using diffusion tensor imaging (DTI). These studies have investigated white matter mostly in terms of tensor metrics such as fractional anisotropy and/or mean diffusivity, and assessed how they may be altered in, or associated with WMH (de Groot et al., 2000; Firbank et al., 2003; Taylor et al., 2007; Vernooij et al., 2008; Lee et al., 2009; Topakian et al., 2009; Altamura et al., 2016; Seiler et al., 2018). Studies commonly report white matter microstructure, as measured using these DTI-derived metrics, to be altered not only within WMH (Bastin et al., 2009; Maniega et al., 2015), but additionally altered within NAWM in patients with such lesions (Firbank et al., 2003; Vernooij et al., 2008; Maniega et al., 2015). As such, DTI has been suggested to be a more sensitive model to assess subtle white matter damage than WMH detected on FLAIR (Charlton et al., 2010; Maillard et al., 2013). However, the diffusion tensor is a limited model that cannot adequately describe white matter microstructure in voxels containing more than one fibre population (i.e., most white matter voxels) (Jeurissen et al., 2013; Jones et al., 2013), and changes to DTI metrics may be difficult to interpret in brain injury (Budde et al., 2011). As such, while useful in detecting abnormalities within white matter, any differences quantified by tensor-based metrics are quite non-specific and in many cases, can be inaccurate or misleading. Consequently, DTI is unable to provide clear indication of the specific nature of microstructural damage within WMH.

In the present work, we utilised single-shell 3-tissue constrained spherical deconvolution (SS3T-CSD) to model white matter microstructure (Dhollander and Connelly, 2016a; Dhollander et al., 2016). There are a number of advantages to this method that enable us to identify particular changes to white matter structures. SS3T-CSD is able to more appropriately model white matter in voxels that also contain other tissue types, by also modelling different tissue compartments in addition to white matter, as has been done for multi-shell data (Jeurissen et al., 2014). The added advantage of SS3T-CSD is that it can model 3 tissue compartments using *single-shell* data alone, enabling both a shorter acquisition time and the ability to investigate historical single-shell data.

In this study, we took into account joint changes to the complete 3-tissue composition: T_W_, T_G_, and T_C_. This enabled us to characterise microstructural properties of tissue when it deviated from that of normal white matter. By characterising WMH in terms of these three signal fractions, we could interpret the resulting 3-tissue profile based on how alike the diffusion signal properties were to those derived from normal white matter, grey matter and CSF. However, careful consideration is required when interpreting the results, based on the context within which we observe changes to the 3-tissue composition. That is, we should not simply interpret increases or decreases in each signal fraction a reflection of alterations in the amount of healthy tissue type from which the response function was derived. For example, an increase in T_G_, as was evident across WMH in comparison to NAWM, should not be interpreted as an increase in actual grey matter, but a shift toward something that has similar diffusion characteristics. This increase in T_G_ could be compatible with astrogliosis, as proliferation of glial cells is known to be a characteristic of WMH, and such a change would likely have a similar effect as grey matter on the diffusion signal: diffusion would still be relatively hindered, but much more isotropic than the healthy white matter represented by T_W_. An increase in the T_C_ fraction could be interpreted as an increase in free fluid, which would likely reflect increases in interstitial fluid that may accompany nearby ependymal breakdown or local myelin or axonal loss (Dhollander et al., 2017).

Importantly, the major advantage of characterising microstructure using this 3-tissue model, is that we could identify heterogeneity within WMH that could not be identified with other imaging modalities *in vivo*. This enabled identification of clear variability across different classes of lesions, and even within confluent lesions that appeared otherwise homogeneous on FLAIR. Such a finding has major implications when investigating these lesions *in vivo*: on the evidence of a technique that enables identification of *in vivo* variability within lesions, it would appear inappropriate to amalgamate all WMH together as a singular pathological entity.

### 4.4 Limitations and future directions

This work represents a preliminary investigation into the *in vivo* heterogeneity of WMH that can be explored with diffusion MRI, and as such there are a number of limitations to our work that should be highlighted. Firstly, given that we do not have histological data to compare our *in vivo* metrics to, we cannot directly interpret the results in terms of their pathological basis. As such, future work that correlates these diffusion metrics with post-mortem histopathology could provide more direct evidence for the specific histological underpinnings of these metrics, and enable broader *in vivo* investigation of disease-relevant pathology within WMH in large cohort studies.

Secondly, the aim of this study was to determine if we could probe *in vivo* variability within WMH using diffusion MRI, rather than to provide a method that could classify or differentiate lesion areas based on these diffusional properties. Future work that characterises lesions based on their specific microstructural properties, rather than on arbitrary visually-guided or purely distance-based schemes could be highly useful, as it could enable the development of novel classification schemes with more meaningful microstructural basis. Such diffusion-based classification schemes could prove particularly valuable when investigating the association of WMH with disease-specific changes, given that certain microstructural changes are likely to be more detrimental than others.

Finally, while we explored the microstructural properties of WMH in healthy elderly individuals and Alzheimer’s disease subjects, in this work, we did not focus on aspects specific to the microstructural characteristics of WMH in Alzheimer’s disease. To this end, the development of the aforementioned, diffusion-based classification scheme in future work could be highly insightful, as it may enable us to identify the microstructural properties of more pathologically harmful WMH and determine and how these might be related to Alzheimer’s disease.

### 4.5 Conclusion

In this study, we were able to detect microstructural heterogeneity within lesions *in vivo*, and identify variability within lesion classes based on their microstructural features through the application of diffusion MRI, and in particular SS3T-CSD. Diffusion MRI is likely more sensitive to underlying pathological features than FLAIR MRI, and would thus be a highly valuable probe to investigate WMH, potentially in conjunction with FLAIR. Given that Alzheimer’s disease subjects have higher lesion load of certain classes and locations of WMH, it would be useful to investigate the particular *in vivo* features that are closely related with disease progression. Future work investigating the microstructural properties of WMH and their clinical and pathological correlates is likely to be highly insightful.

## Abbreviations

AIBL: Australian Imaging Biomarkers and Lifestyle study of ageing
CSD: Constrained spherical deconvolution
CSF: cerebrospinal fluid
DTI: Diffusion tensor imaging
DWI: Diffusion-weighted imaging
FLAIR: Fluid-attenuated inversion recovery
FOD: Fibre orientation distribution
GM: grey matter
HIST: HyperIntensity Segmentation Tool
NAWM: Normal-appearing white matter
SS3T-CSD: Single-shell 3-tissue constrained spherical deconvolution
T_C_: Cerebrospinal fluid-like signal fraction
T_G_: Grey matter-like signal fraction
T_W_: White matter-like signal fraction
WM: white matter
WMH: White matter hyperintensities

## Acknowledgements

We thank the patients, researchers and clinicians involved in the Australian Imaging, Biomarkers, and Lifestyle study of ageing. The Florey Institute of Neuroscience and Mental Health acknowledges the strong support from the Victorian Government and in particular the funding from the Operational Infrastructure Support Grant. The authors acknowledge the facilities and scientific and technical assistance of the National Imaging Facility, a National Collaborative Research Infrastructure Strategy (NCRIS) capability, at the Florey Institute of Neuroscience and Mental Health.

