## Supplementary Material for "*In vivo* microstructural heterogeneity of white matter lesions in Alzheimer’s disease using tissue compositional analysis of diffusion MRI data"

**Supplementary Table 1: WMH volume comparisons between amyloid-positive and amyloid-negative healthy control subjects**

| | Amyloid-negative HC (n = 63) | Amyloid-positive HC (n = 31) | Statistics | *P*-value |
|---|---|---|---|---|
| **Regional WMH volumes ($cm^3$), mean (SD)** | | | | |
| **Periventricular** | 7.04 (7.69) | 8.56 (10.87) | | |
| **Deep** | 0.81 (0.76) | 1.04 (1.71) | | |
| **ROI 1** | 2.17 (1.77) | 2.63 (2.32) | | |
| **ROI 2** | 2.42 (2.64) | 2.68 (2.45) | | |
| **ROI 3** | 3.23 (4.27) | 4.24 (7.70) | | |
| **Cubic root WMH volumes, mean (SD)** | | | | |
| **Total WMH** | 1.79 (0.62) | 1.90 (0.66) | $F(1, 90) = 0.07$ | 0.79 |
| **Periventricular WMH** | 1.70 (0.63) | 1.82 (0.66) | $F(1, 90) = 0.08$ | 0.78 |
| **Deep WMH** | 0.84 (0.30) | 0.83 (0.42) | $F(1, 90) = 0.07$ | 0.79 |
| **ROI 1 WMH** | 1.20 (0.36) | 1.26 (0.42) | $F(1, 90) = 0.16$ | 0.69 |
| **ROI 2 WMH** | 1.17 (0.48) | 1.25 (0.45) | $F(1, 90) = 0.07$ | 0.79 |
| **ROI 3 WMH** | 1.23 (0.59) | 1.32 (0.65) | $F(1, 90) = 0.01$ | 0.92 |

**Supplementary Material: Lobar analysis**

The anatomical location of the WMH was considered for a third analysis. WMH voxels were classified according to the brain lobe in which they were located. The frontal, parietal, occipital and temporal lobes were manually delineated on the population template image, by extrapolating surface landmarks of their boundaries deeper into the white matter at each axial slice. WMH voxels were classified into frontal, parietal, occipital, or temporal, based on which lobe they overlapped with.

*Regional white matter hyperintensity volumes*

WMH was significantly higher in Alzheimer's disease patients compared to controls in all lobes (frontal: $F(138) = 11.25$, $p = 0.001$; occipital: $F(138) = 19.11$, $p < 0.001$; parietal: $F(138) = 5.50$, $p = 0.02$; temporal: $F(138) = 4.05$, $p = 0.046$).

*Lobar regional analysis of 3-tissue profile*

In order to determine if there was a significant difference in the 3-tissue profile of WMH in different lobar locations, we performed an analysis to compare between WMH within frontal, parietal, occipital and temporal white matter. Supplementary Figure 1 shows the ternary plot exhibiting the profiles of WMH within each lobe, with the corresponding log-ratio transformed plot shown in Supplementary Figure 2. Multivariate analyses showed statistically significant differences between NAWM and WMH is all regions (frontal WMH: $F(2, 281) = 7000.12$, $p < 0.001$, Pillai's trace = 0.980; occipital WMH: $F(2, 279) = 3257.86$, $p < 0.001$, Pillai's trace = 0.959; parietal WMH: $F(2, 276) = 3334.89$, $p < 0.001$, Pillai's trace = 0.960; temporal WMH: $F(2, 268) = 3617.84$, $p < 0.001$, Pillai's trace = 0.964). There was a significant difference in the diffusional profile between frontal WMH and WMH in all other regions (occipital: $F(2, 279) = 338.72$, $p < 0.001$, Pillai's trace = 0.708; parietal: $F(2, 276) = 182.45$, $p < 0.001$, Pillai's trace = 0.569; temporal: $F(2, 268) = 259.29$, $p < 0.001$, Pillai's trace = 0.659), as well as between occipital and parietal ($F(2, 274) = 7.18$, $p = 0.001$, Pillai's trace = 0.050), occipital and temporal ($F(2, 266) = 34.72$, $p < 0.001$, Pillai's trace = 0.207), and temporal and parietal ($F(2, 263) =$

23.49, $p < 0.001$, Pillai's trace = 0.152), all with a Bonferroni-corrected significant p-value of 0.005.

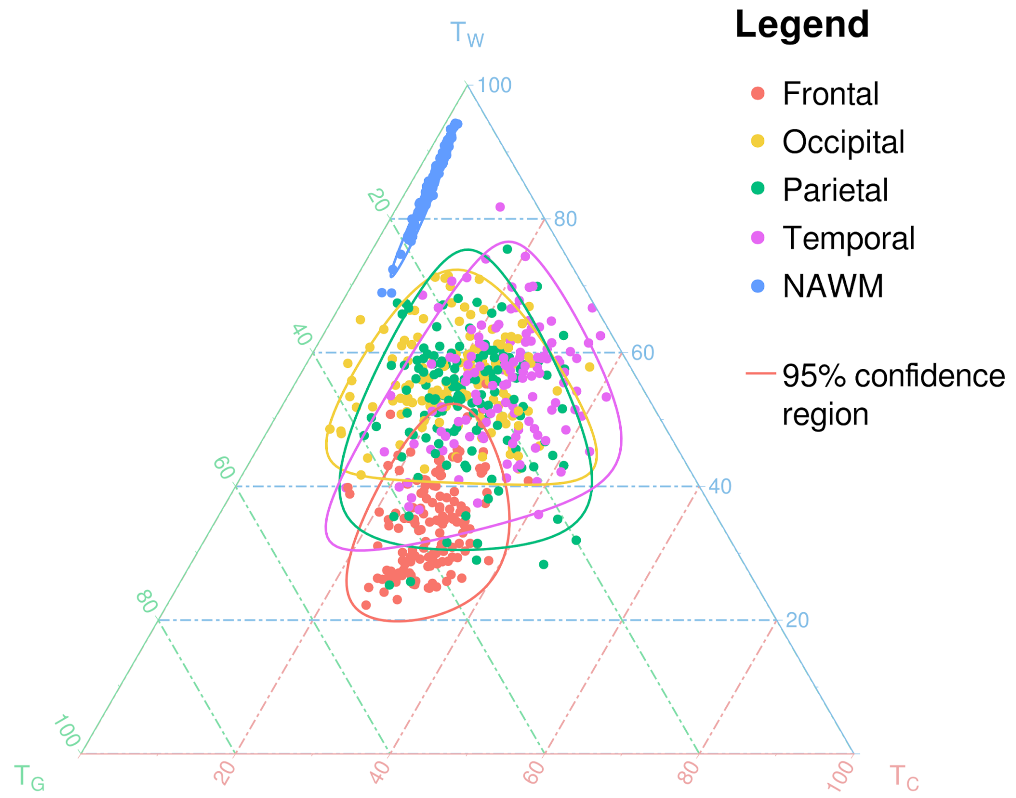


**Supplementary Figure 1: Ternary plot showing diffusional profile of lesions in each lobar region.** The relative $T_W$, $T_G$ and $T_C$ fraction was computed for WMH in each lobar region, and within NAWM. The mean $T_W$, $T_G$ and $T_C$ fractions was computed for all lesional voxels within each lobar region, for each subject. The mean profiles for each lobar region is plotted for all Alzheimer's disease and healthy elderly subjects on the ternary plot. Unlike the previous analyses, there was no clear fraction that could differentiate WMH within each lobe from one another, but the mean diffusional profile for each lobar region could be distinguished from all other regions, and from NAWM, when statistical analyses were performed on the isometric log ratio transformed data.

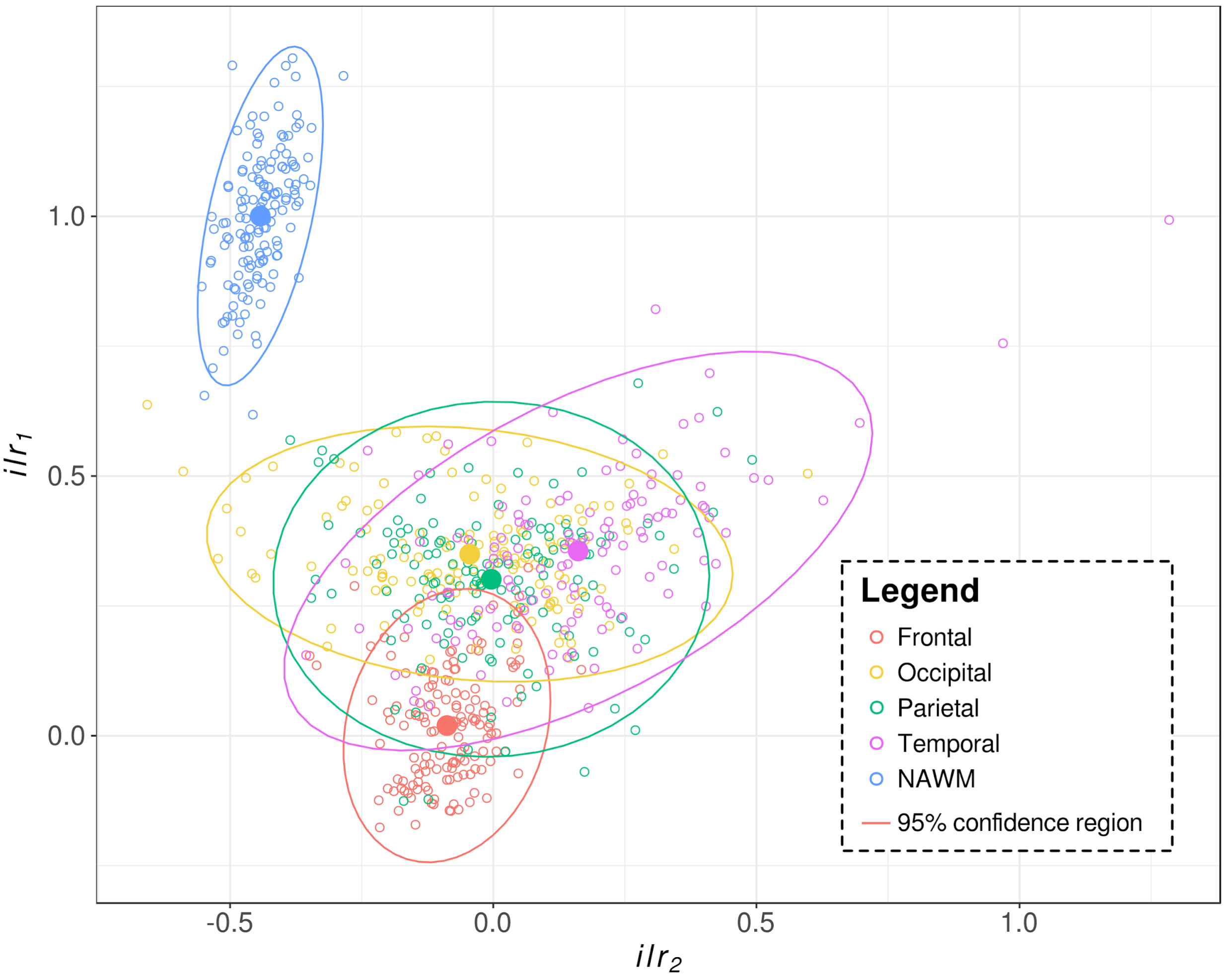


**Supplementary Figure 2: Isometric log-ratio transformed diffusional data comparing WMH in each region and NAWM.** As shown and described in Fig. 5 and in Supplementary Fig. 1, an isometric log-ratio transform was applied to the relative signal fractions ($T_W$, $T_G$, $T_C$) to transform the compositional data, in this case for the regional analysis. Data points reflect the transformed diffusional profile from a single subject for WMH in the frontal (red), parietal (green), occipital (yellow), or temporal (purple) lobe, or for normal-appearing white matter (NAWM; blue). The centroid for each region or NAWM is shown as a solid circle, while the population 95% confidence ellipses is shown as a solid line.
